# A combinatorial genetics approach reveals limits to redundancy within *Plasmodium falciparum* invasion ligand families

**DOI:** 10.64898/2026.02.23.706623

**Authors:** Eleanor Silvester, Carla Briggs, Alazhar Colombowala, Anna Kuroshchenkova, Victor Flores, Emma Kals, Julian C. Rayner

## Abstract

To maintain infection in the human bloodstream, *Plasmodium falciparum* parasites undergo repeated cycles of invasion of and multiplication inside red blood cells (RBCs). Two protein families, the Erythrocyte binding-like (EBA) and Reticulocyte binding-like (Rh) proteins, are known to play a key role in invasion, mediating early stages of attachment of the *P. falciparum* merozoite to the host RBC. There is a degree of redundancy within these families, such that disrupting the function of individual EBA/Rh proteins *in vitro* is not sufficient to prevent invasion. By employing a novel approach to disrupt multiple EBA and Rh genes in combination, we systematically assessed functional interdependency across these families for the first time. This analysis, and further characterisation of mutant parasites, revealed that disruption of some pairs of EBA/Rh ligands significantly impacted *P. falciparum* invasion, whereas others did not. Disruption of PfEBA175 in combination with either PfRh2b or PfRh4 significantly reduced parasite multiplication *in vitro*, indicating that the remaining EBA/Rh proteins could not fully compensate for the absence of these three key invasion ligands. PfEBA175, PfRh2b and PfRh4 may therefore represent a critical nexus of the parasite invasion machinery, and combinatorial approaches that target these specific ligands could be a beneficial therapeutic approach.

## Introduction

There were an estimated 263 million cases of malaria globally in 2023, causing an estimated 597,000 deaths ^1^, with most deaths caused by *Plasmodium falciparum* parasites. Following transmission by an infected *Anopheles* mosquito, the parasite transitions through a developmental stage within the liver, before establishing repeated cycles of replication within human red blood cells (RBCs) to maintain infection. It is the blood stage of infection, which features parasite sequestration in blood vessels to prevent splenic clearance, that causes morbidity and mortality. During the blood stage, merozoites egress from an infected RBC before rapidly invading neighbouring ones ^2^. In the brief period where merozoites are extracellular, proteins on their surface are exposed to the immune system, and antibodies to merozoite proteins are a major component of the immune response to *Plasmodium falciparum* ^3^.

*Plasmodium falciparum* parasites employ multiple evasion mechanisms to prevent antibody-mediated targeting of their invasion machinery. One mechanism is to limit exposure to the immune system. The essential and highly conserved invasion protein PfRh5 is an example as it has yielded promising results in early stage vaccine trials, but is known to be poorly immunogenic in natural infections, limiting the potential for natural boosting ^4,5^. An alternative mechanism is generating multiple distinct variants of an antigen, limiting antibody cross-reactivity. The critical invasion protein PfAMA1 is highly polymorphic^6^, preventing the development of a strain transcending protective response and reducing efficacy in vaccine trials^7,8^. A third mechanism is redundancy in important invasion ligands^9,10^. The Erythrocyte binding-like (EBA) and Reticulocyte binding-like (Rh) proteins are encoded by two distinct multigene families, but with the exception of PfRh5 (which has a very different structure to other Rh proteins), all are hypothesised to fulfil a similar and interchangeable role during invasion. Members of these protein families are released from apical organelles onto the merozoite surface and engage with receptors on the RBC surface, this tight interaction forms a critical step early in the invasion process. There are 4 EBA proteins (PfEBA140, PfEBA175, PfEBA181 and PfEBL1), and 4 Rh proteins (PfRh1, PfRh2a, PfRh2b, PfRh4) expected to function in at least some strains of *P. falciparum*. Experimental genetics has shown that it is possible to disrupt expression of any of these eight EBA and Rh genes without serious detriment to the parasite^11–14^ and expression of EBA and Rh genes has been observed to vary in different *P. falciparum* strains^15,16^. This redundancy is thought to aid immune evasion, as well as allow adaptation to variation in RBC receptors that may be encountered *in vivo*.

However, the true extent of redundancy and interchangeability between EBAs and Rhs has never been clearly experimentally established, and all *Plasmodium* genomes sequenced to date appear to retain at least one EBA and one Rh orthologue^17^. In the zoonotic species *Plasmodium knowlesi*, where only one EBA and one Rh orthologue can function in invasion of human RBCs, conditional disruption approaches have revealed distinct roles during invasion for these two families ^18^. In *P. falciparum,* unique roles for the EBA and Rh proteins are more complicated to disentangle because of the larger number of family members, and there is clearly some level of redundancy, as single gene mutants are not significantly impaired. Even here however there is evidence for some interdependence and interaction between members of these families. For example, increased expression of PfRh4 was observed following disruption of PfEBA175 ^11,13^. PfEBA175 disruption was also associated with increased sensitivity to antibodies targeting Rh2 proteins ^11^. However, a family-wide assessment of functional interdependency between the Rh and EBA proteins had not previously been undertaken.

We sought to systematically test the viability of all possible combinations of EBA and Rh null mutants and thereby identify pairs of genes which when disrupted together were more detrimental to parasite survival. To do this at higher throughput we developed a pooled transfection approach which utilised next generation sequencing to assess viability before generating individual double, and in some cases triple, mutant lines using traditional approaches. In doing so, we revealed that it was possible to disrupt any two EBA or Rh genes and retain viability, but that some combinations had more detrimental effects. Subsequent detailed phenotypic analysis using both genetic and antibody-mediated approaches highlighted PfRh2b, PfRh4 and PfEBA175 as together playing key roles in invasion in the 3D7 strain. Invasion assays indicated that these ligands were involved in compensating for each other’s absence, and disruption of either PfRh2b or PfRh4 in combination with PfEBA175 was more detrimental to growth in both static and shaking conditions. The lack of compensation by the remaining EBA and Rh ligands suggests that not all family members are truly functionally equivalent, raising the possibility that targeting only a specific subset of ligands could form part of an effective intervention strategy.

## Results

### A novel strategy to disrupt the function of multiple EBA and Rh invasion ligands in combination

The EBA and Rh invasion ligands are Type 1 membrane proteins that have a C-terminal transmembrane domain followed by a short cytoplasmic domain. The domain required for binding to the host erythrocyte is located in the N-terminal region of the large extracellular domain, immediately following the signal sequence ^19–25^. Disruption of this region is therefore expected to prevent EBA or Rh proteins from binding their target receptors and hence functioning during erythrocyte invasion. We used CRISPR/Cas9 guide RNA (gRNA) targeting to introduce 4 frameshift insertions and 6 extra stop codons into the binding region of each EBA/Rh gene, to prevent expression of a full-length functional protein. In the case of PfRh2a and PfRh2b, which share an identical binding region, the disruption was instead targeted to unique regions in the C-terminus, the only region that differs in these adjacent genes (Figure 1A). This method was initially used to generate *P. falciparum* cell lines with a single EBA or Rh gene disrupted (EBA/Rh single knockout). All EBA and Rh ligands expected to have functionality in the 3D7 strain used were targeted (except for PfRh5, which as noted above has a functionally distinct role). PfEBA165 and PfRh3, which have 5’ frameshift mutations in all known *P. falciparum* isolates^26–28^, and PfEBL1 which has a 5’ frameshift mutation in some *P. falciparum* strains including 3D7^29^ were excluded from the strategy. Disruption of Pfs230 (PF3D7_0209000) was included as a control as it is not expected to function in the asexual blood stage of the life cycle. Pfs230 is expressed in gametocytes and is important for the interaction between male gametes and RBCs in the mosquito midgut to form exflagellation centres, a critical step for successful onward transmission ^30^. Integration of the frameshift mutations at the targeted loci was confirmed by PCR (Supplementary figure 1). To validate the method, disruption of the PfEBA175 protein by frameshift insertion was confirmed using an antibody targeting a region downstream of the inserted mutations (Figure 1B).

**Figure 1.**
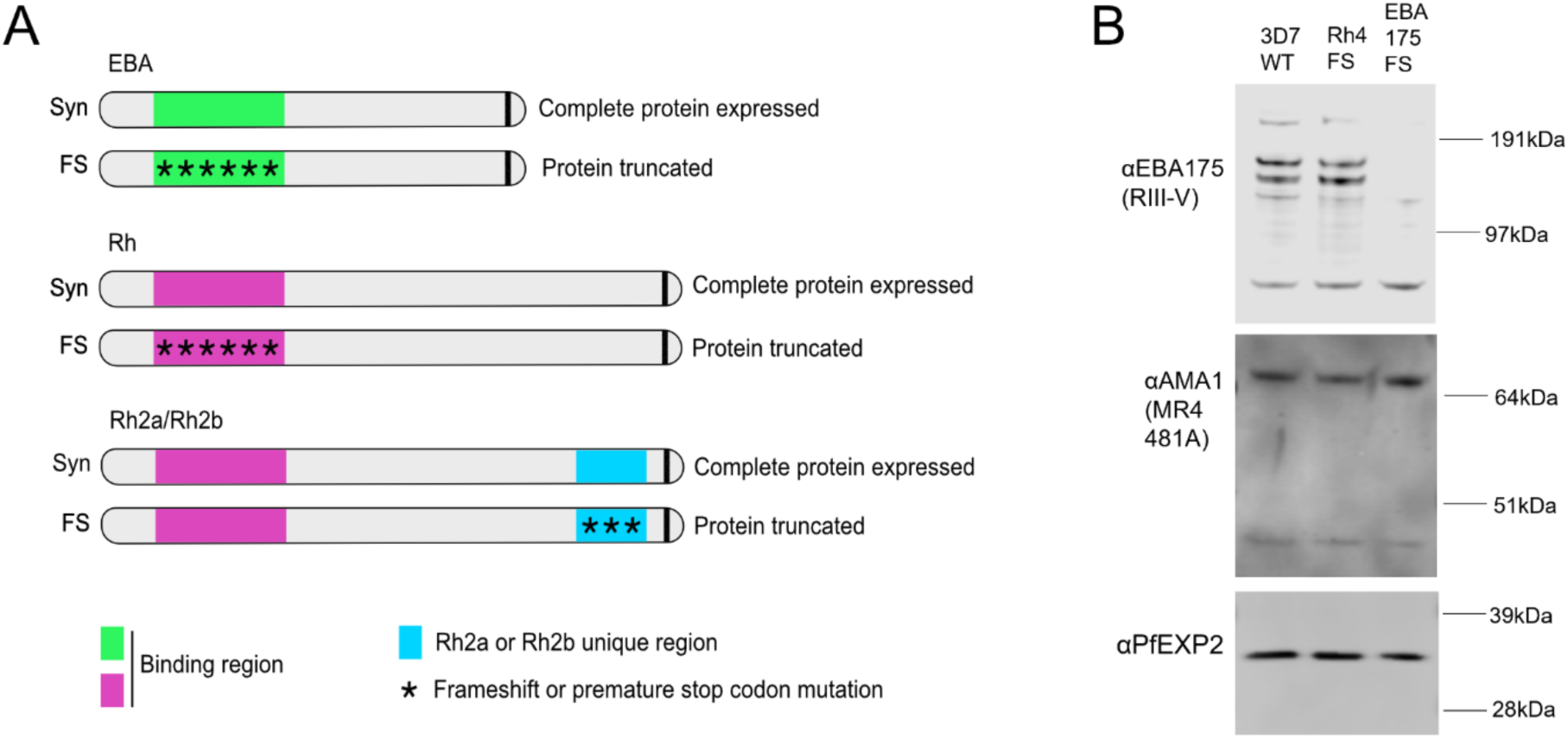
Strategy to disrupt EBA/Rh genes. A) Target genes are edited by introduction of a 300-500bp recoded nucleotide sequence that does not alter the amino acid sequence (Synonymous, Syn) or a recoded sequence that introduces frameshift mutations and premature stop codons (FS). These sequences are targeted to the region of the gene important for erythrocyte binding, except in the case of PfRh2a and PfRh2b, which are identical across most of the coding sequence, so mutations were targeted to the unique region at the C-terminus. B) Western blot confirming disruption of PfEBA175 protein expression by introduction of frameshift mutations.

Single knockout parasite lines with disruption targeting PfEBA175, PfEBA181, PfEBA140, PfRh1, PfRh2a, PfRh2b or PfRh4 were all viable, as expected based on data from previous studies^11–14^. Targeting multiple EBA and Rh ligands in combination could have a greater impact on invasion of RBCs and viability. We therefore aimed to systematically generate double knockout parasites with all possible combinations of EBA and Rh genes disrupted. To perform such an all vs all screen for the first time, we developed a pooled transfection approach for higher throughput and used Next Generation sequencing to identify recoverable double knockout combinations. The CRISPR/Cas9 editing efficiency in *P. falciparum* is variable and can be affected by gRNA selection. We therefore first identified the most effective gRNAs for each target gene during the generation of single knockout lines and used these gRNAs in the pooled transfections.

EBA/Rh single knockout parasite lines were transfected with Cas9/gRNA plasmids along with two possible repair templates for each gene. One repair template introduced frameshift mutations into the target gene (as used to generate the single knockouts), and the other repair template introduced a synonymous recodonised sequence which targeted the same region but did not disrupt the protein coding sequence (Figure 2A). The synonymous sequence was included as a control to account for technical variability in *P. falciparum* transfections. The synonymous and frameshift recodonised repair sequences differ only in the presence of the frameshift mutations and a unique 30 base pair sequence for easier detection of frameshift edits. Following transfection, the potentially edited region of each target gene was amplified using primers flanking the homologous repair region, meaning that only edits that had been incorporated into the genome were amplified, and the same primers would amplify both frameshift and synonymous edited regions. The sequences of the amplified fragments were confirmed using next-generation sequencing, yielding on average 2.8 million read pairs, each amplicon was sequenced in its entirety by most of the Illumina libraries, with an average depth of coverage of at least 1 million reads per nucleotide (Supplementary table 1). The relative recovery of frameshift and synonymous edits in the sequence data was used to indicate cases where a frameshift edit could have an impact on parasite viability. Simplistically, if both the synonymous and frameshift edits were recovered this would indicate that the double knockout combination tested was viable, whereas if only the synonymous edit was detected this would indicate a double knockout combination that was detrimental to the parasite.

**Figure 2.**
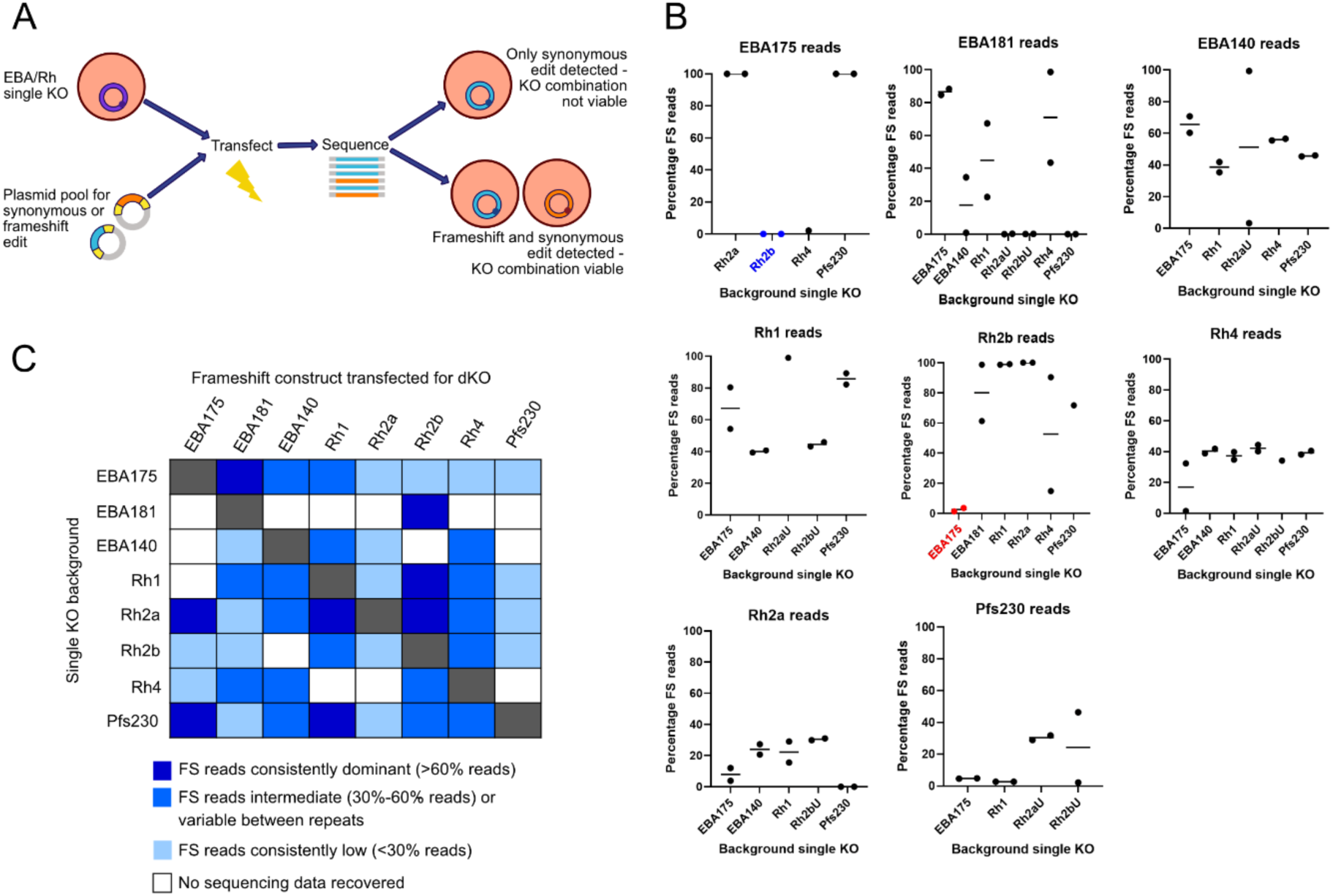
A pooled transfection and sequencing strategy reveals that most EBA/Rh double knockout combinations are viable and identifies combinations that may be detrimental. A) Pooled strategy to identify viable EBA and Rh double knockout (KO) combinations. B) Next generation sequencing was used to detect synonymous and frameshift edits in targeted EBA and Rh genes. The percentage of total reads that correspond to a frameshift (FS) edit are plotted for each gene on a separate graph, with the transfected cell lines these edits were detected in indicated on the x axis. PfEBA175 FS edits in the PfRh2b single KO background are highlighted in blue. PfRh2b FS edits in the PfEBA175 single KO background are highlighted in red. C) Summary of pooled sequencing results with the proportion of frameshift reads recovered for each combination of genes indicated by different shades of blue.

In all cases where synonymous edits were recovered for a target gene, there was also recovery of frameshift edits (Supplementary figure 2, Supplementary table 2). The collection of edits detected within the pools indicate that most EBA and Rh double knockout combinations are viable and significantly extends previous studies of redundancy and interchangeability within and between these gene families. In some cases, however, the number of reads recovered for a frameshift edit were very low (Figure 2B), for example recovery of PfEBA175 frameshift reads in the PfRh2b single knockout background (sample 1 - 0.001% of reads, sample 2 – 0.00005% of reads). This was despite good recovery of PfEBA175 frameshift reads in the PfRh2a and Pfs230 control single knockout backgrounds, where the same PfEBA175 targeting guide and PfEBA175 repair templates were used. Similarly, the number of PfRh2b frameshift reads recovered in the PfEBA175 single knockout background was much lower than the number of synonymous reads recovered (sample 1 frameshift – 1.3% of reads, sample 2 frameshift - 3.6% of reads). In contrast, the number of PfRh2b frameshift reads recovered was higher than the number of synonymous reads in the PfEBA181, PfRh1 and Pfs230 single knockout backgrounds, where again the same PfRh2b targeting guide and PfRh2b repair templates were used. These results together indicated that the parasite may not be able to readily compensate for the combined absence of PfRh2b and PfEBA175, and invasion may be less efficient in double knockout parasites that lack these two ligands. There was also some evidence that the PfRh4 and PfEBA175 double knockout combination may be detrimental, though in this case, recovery of frameshift reads was not as consistently low as the PfRh2b/PfEBA175 combination.

The only double disruption combination that was not recovered from the pools was PfRh2b+PfEBA140, but this was because we were unable to generate amplicons for sequencing. There was no recovery of parasites from the pool containing the PfRh2b edit in the PfEBA140 single knockout background, and in the PfRh2b single knockout background it was not possible to amplify the region of PfEBA140 targeted by the edit. However, we were subsequently able to generate the PfEBA140+PfRh2b double mutant in an independent transfection, confirming the viability of this double mutant combination.

Previous studies have produced and characterised a limited number of EBA double mutants (PfEBA175/181, PfEBA175/140) ^11^ as well as a PfRh2a/Rh2b double mutant^31^. We have significantly expanded the number of viable EBA/Rh double mutants, detecting 20 unique combinations in our pooled transfections (Figure 2C). This includes generation of cross family mutant combinations for the first time, combining disruption of one EBA and one Rh gene.

### Characterisation of EBA/Rh double knockout lines in static and shaking growth conditions

To validate the pooled transfection data, EBA/Rh double knock out parasite lines were generated by individually transfecting single knock out backgrounds with frameshift repair constructs. 14 unique EBA/Rh double mutants were generated and cloned by limiting dilution to produce clonal lines for which growth and invasion characteristics could be analysed in greater depth (Supplementary Figure 3). We characterised growth in both static and shaking conditions (Figures 3A and 3B), as shaking conditions may be more reflective of the dynamic environment in which invasion takes place in a human host. We used shaking conditions that correspond to the point at which the blood transitions from sedimentation into suspension, where modelling predicts exerted forces comparable to those that may be experienced in the microvasculature. These conditions had previously revealed growth and invasion phenotypes for EBA/Rh single knockout lines not seen in static assays ^32^.

**Figure 3.**
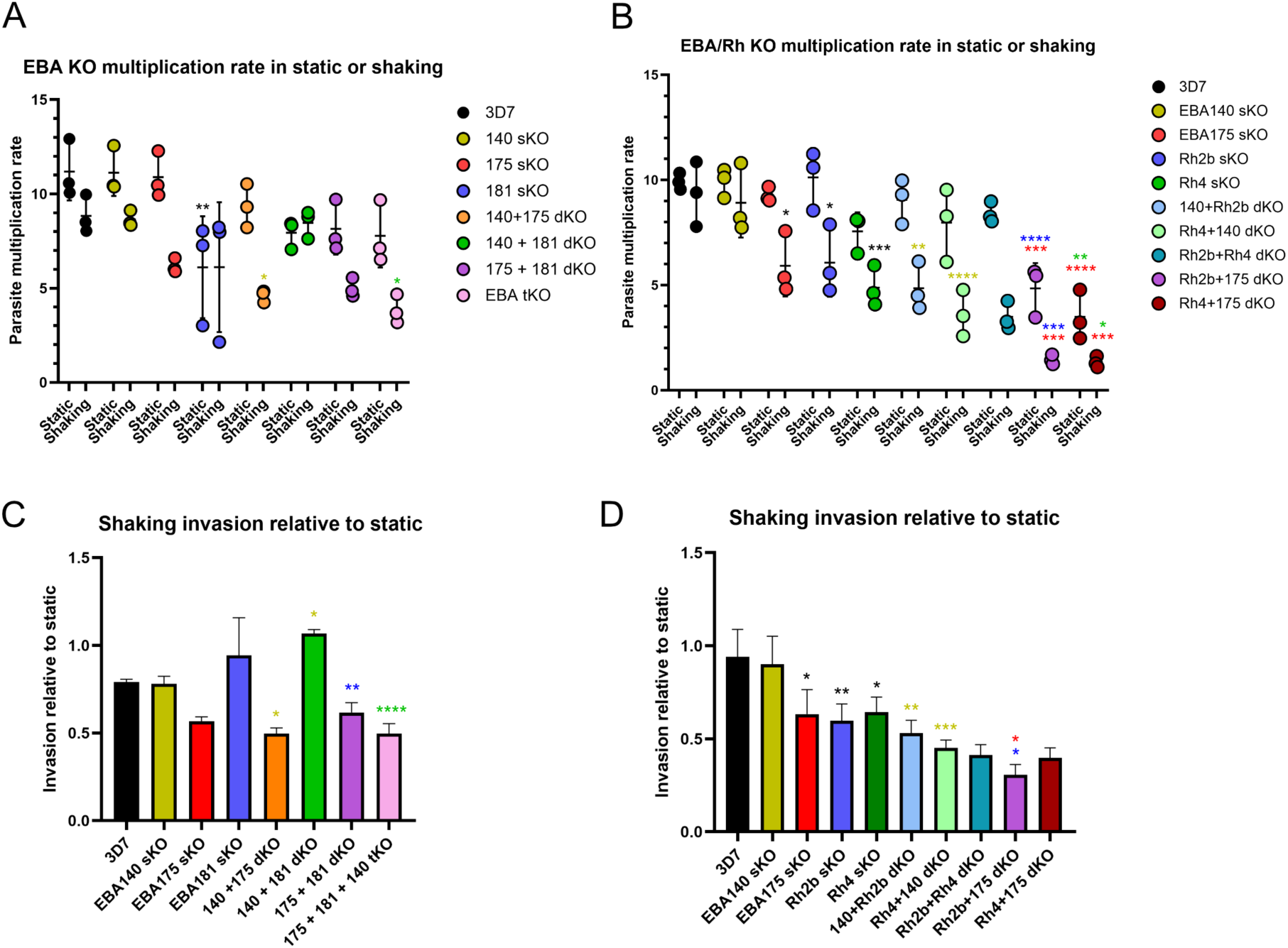
Disruption of certain EBA and Rh genes reduces proliferation in static and/or shaking conditions. A) Multiplication rate of EBA single, double or triple knockout lines in static or shaking conditions. B) Multiplication rate of EBA/Rh single and double knockout lines in static or shaking conditions. C) Parasite multiplication over 48h in shaking conditions relative to static conditions for EBA single, double and triple knockout cell lines. D) Parasite multiplication over 48h in shaking conditions relative to static conditions for PfEBA175, PfEBA140, PfRh2b and PfRh4 single and double knockout cell lines. (*P<0.05, **P<0.01, ***P<0.001, ****P<0.0001, two-way analysis of variance with Tukey’s multiple comparisons test). The error bars represent the mean ± s.d of three biological replicates. Statistics are shown for comparisons between single knockout lines and 3D7 or double knockout cell lines and their corresponding single knockout lines (* are coloured to match the comparator line). Comparisons that did not pass the significance threshold (P<0.05) are not plotted.

In shaking conditions, PfEBA175, PfRh2b and PfRh4 single knockout cell lines demonstrated significantly reduced proliferation compared to the parental 3D7 strain. In contrast, the PfEBA140 single knockout cell line behaved like the parental 3D7 strain, indicating that certain invasion ligands are more important than others in maintaining productive invasion under shaking conditions. This could relate to the structure or abundance of the ligands and receptors involved, or to the affinity of their interactions. Interestingly, the PfEBA181 single knockout cell line was less impacted by shaking than the parental 3D7 strain. These data were largely consistent with previous observations of single EBA and Rh knockouts in the NF54 strain ^32^.

Testing EBA double knockout combinations in shaking conditions revealed a hierarchy where the effect caused by disruption of one EBA was dominant (Figure 3C). Disruption of PfEBA181 reduced sensitivity to shaking relative to the 3D7 strain, and this less sensitive phenotype was also observed in the PfEBA140+PfEBA181 double knockout line. The enhanced sensitivity to shaking associated with disruption of PfEBA175 was observed in both the PfEBA140+PfEBA175 and PfEBA175+PfEBA181 double knockout lines, revealing the dominant effect of targeting PfEBA175. The PfEBA175+PfEBA181 double knockout line was further modified to introduce frameshift mutations into PfEBA140, yielding a viable EBA triple knockout cell line for the first time. The 3D7 strain used to generate the cell lines in this study had historically acquired frameshift mutations in PfEBL1^29^, therefore PfEBA175, PfEBA140 and PfEBA181 are expected to be the only functional EBA ligands remaining in this strain. The continued presence of PfEBL1 frameshift mutations in the EBA triple knockout was confirmed by Sanger sequencing (Supplementary figure 4), this line is therefore not expected to express any functional EBA proteins. Surprisingly, the absence of any functional EBA proteins was relatively well tolerated, with the parasite multiplication rate of the triple knockout comparable to that of the parent PfEBA175+PfEBA181 double knockout in both static and shaking conditions.

We next assessed how disrupting one invasion ligand from each of the two protein families influenced growth in static and shaking conditions, compared to disruption of two ligands from within the same family. Double knockout combinations tested featured PfEBA175, PfEBA140, PfRh2b and PfRh4. Multiple attempts were required before PfRh2b+PfEBA175 and the PfRh4+PfEBA175 double knockout parasite lines could be generated and the multiplication rate of these cell lines in static conditions is significantly lower than their parent single knockout lines (Figure 3B). These data are consistent with the poor recovery of frameshift edits for these combinations in the pooled transfections and suggest that the combination of either PfRh4 or PfRh2b with PfEBA175 is particularly detrimental in 3D7 parasites. The proliferation of PfRh2b+PfEBA175 and PfRh4+PfEBA175 double knockout lines is reduced even further in shaking, and in the case of PfRh2b+PfEBA175 the reduction caused by shaking is significantly more than when either gene is targeted alone (Figure 3D). The proliferation of the PfRh2b+PfRh4 double knock out cell line is also dramatically reduced in shaking conditions, but static growth is comparable to the parent single knockout lines.

These results indicate that targeting an Rh and an EBA ligand in combination is more disruptive to invasion in static and shaking conditions than targeting two EBA or two Rh ligands. Furthermore, there is clearly not complete functional equivalence within families, as the absence of a functional PfEBA175 is not compensated by the remaining EBA proteins. Critically, the greatest impact on invasion is observed when either PfRh2b or PfRh4 is disrupted in combination with PfEBA175.

### EBA and Rh disruption alters invasion into enzyme treated red blood cells

To further investigate invasion pathway preferences and how mutant parasites compensate for absent invasion ligands, we carried out invasion assays using RBCs that had been pre-treated with enzymes. RBC receptors for EBA/Rh ligands are known to have differing sensitivities to enzyme treatments, so these enzymes can be used to indicate which protein interactions are contributing to invasion by a parasite cell line. Neuraminidase cleaves sialic acid from cell surface proteins and impacts binding of EBA ligands and PfRh1 to their receptors ^14,33–38^. Chymotrypsin is a serine protease shown to inhibit PfRh2b, PfRh4 or PfEBA181 mediated invasion ^12,23,39,40^. Invasion into cell trace far red (CTFR) labelled erythrocytes that had been pre-treated with neuraminidase or chymotrypsin (Figures 4A and 4B, Supplementary figure 5) was assessed for EBA/Rh knockout parasite lines. Disruption of PfRh4 was associated with reduced invasion into neuraminidase treated erythrocytes, with invasion reduced further in an PfRh2b+PfRh4 double knockout cell line. The absence of PfRh2b and PfRh4 therefore forces parasites to become more dependent on ligands that engage neuraminidase-sensitive receptors. Disruption of PfRh2b or PfEBA175 individually did not significantly alter neuraminidase sensitivity, however the disruption of PfRh2b and PfEBA175 together resulted in neuraminidase resistant invasion with double knockout parasites invading neuraminidase treated erythrocytes more efficiently than untreated erythrocytes. This suggests the PfRh2b+PfEBA175 double knockout line has become heavily dependent on a neuraminidase-resistant receptor for invasion, possibly the PfRh4 receptor, complement receptor 1. The PfRh4+PfEBA175 double knockout has a neuraminidase sensitivity profile that is intermediate between the PfRh4 and PfEBA175 single knockout profiles. This indicates that the increased neuraminidase sensitivity observed for the PfRh4 single knockout is not only related to an increased dependence on PfEBA175, but also on other neuraminidase sensitive protein interactions.

**Figure 4.**
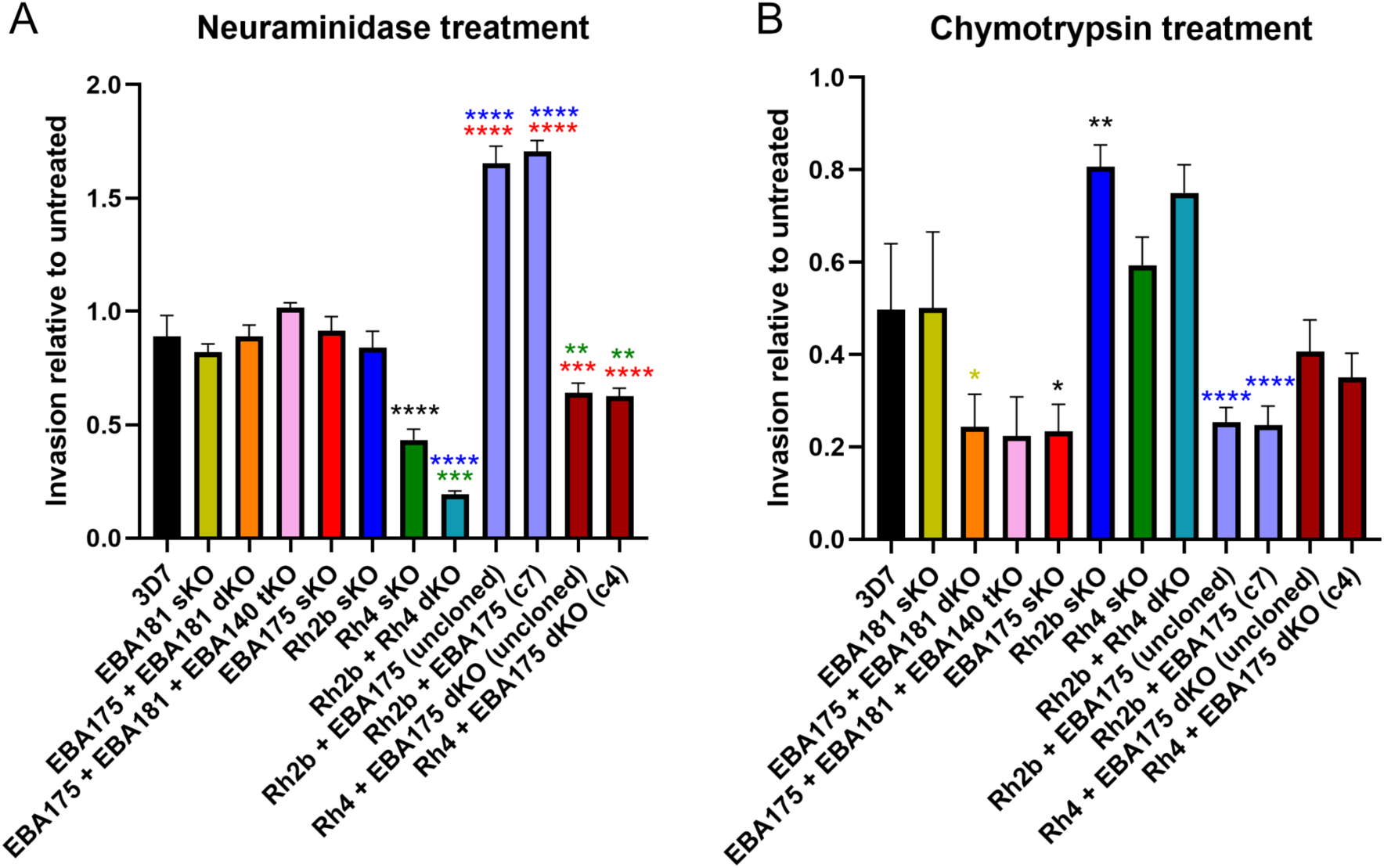
Disruption of certain EBA and Rh genes alters invasion into enzyme treated red blood cells. A) Parasite invasion into neuraminidase treated RBCs relative to invasion into untreated RBCs for EBA/Rh single, double and triple knockout lines. B) Parasite invasion into chymotrypsin treated RBCs relative to invasion into untreated RBCs for EBA/Rh single, double and triple knockout lines (*P<0.05, **P<0.01, ***P<0.001, ****P<0.0001, one-way analysis of variance with Tukey’s multiple comparisons test). The error bars represent the mean ± s.d of three biological replicates. Statistics are shown for comparisons between single knockout lines and 3D7 or double knockout cell lines and their corresponding single knockout lines (* are coloured to match the comparator line). Comparisons that did not pass the significance threshold (P<0.05) are not plotted.

There was a significant negative correlation between the neuraminidase and chymotrypsin sensitivities of the EBA/Rh knockout parasite lines tested, except for the PfRh2b single knockout line which was resistant to both neuraminidase and chymotrypsin treatments (Supplementary figure 6). Disruption of PfEBA175 was associated with increased sensitivity to chymotrypsin treatment. This effect was similar in the EBA double and triple knockout cell lines indicating that PfEBA175 is the dominant EBA and in its absence the parasite does not use the remaining EBA ligands to compensate. The PfRh2b+PfEBA175 double knockout line is also sensitive to chymotrypsin, whereas the PfRh4+PfEBA175 double knockout line is the least chymotrypsin sensitive of the PfEBA175 knockouts. This indicates that in the absence of PfEBA175 and PfRh2b, the parasite may become more reliant on PfRh4 for invasion, but in the absence of both PfRh4 and PfEBA175 there is compensation with multiple alternative ligands (which may include PfRh2b).

The enzyme treatment results support a model where only three ligands (PfEBA175, PfRh2b and PfRh4) from within the Rh and EBA families are major contributors to the parasite invasion phenotype in 3D7. Disruption of PfEBA175, PfRh2b or PfRh4 individually resulted in altered invasion pathway usage as indicated by altered enzyme sensitivities. The additive effect of disrupting PfRh2b and PfRh4 or PfRh2b and PfEBA175 in combination points towards an interdependence between these ligands, where one plays a role in compensating when others are absent.

### Disrupting PfRh2b, PfRh4 and PfEBA175 invasion ligands in combination significantly impairs invasion

To test whether targeting all three critical ligands, PfRh2b, PfRh4 and PfEBA175, in combination caused further disruption, the PfRh2b+PfRh4 double knockout cell line was modified at the PfEBA175 locus to enable inducible knockdown. A 3xHA tag and a glmS ribozyme sequence were introduced after the coding region of PfEBA175 using the selection linked integration method ^41^ (Supplementary figure 7). Addition of glucosamine activates glmS ribozyme self-cleavage destabilising the target transcript and reducing protein expression. Induction with glucosamine to knock down PfEBA175 expression significantly reduced parasite multiplication in two clones compared to uninduced controls (Figure 5A). In contrast, addition of glucosamine did not reduce parasite multiplication in the parental PfRh2b+PfRh4 double knockout cell line. While the effect on parasite multiplication was significant in the PfRh2b+PfRh4 double knockout + PfEBA175 knockdown clones, the reduction in parasite numbers was only 20-40% relative to uninduced controls after 96 hours of treatment. Effective knockdown of PfEBA175 in the PfRh2b+PfRh4 double knockout + PfEBA175 glmS clones following induction was confirmed with both an antibody targeting PfEBA175 and an antibody targeting the HA tag (Figure 5B). Quantification of the western blot indicated that following induction, HA-tagged PfEBA175 levels were reduced by 94% relative to uninduced controls (in both clones tested), but full-length PfEBA175 was still detectable after glucosamine treatment, and may be sufficient to fulfil some level of function.

**Figure 5.**
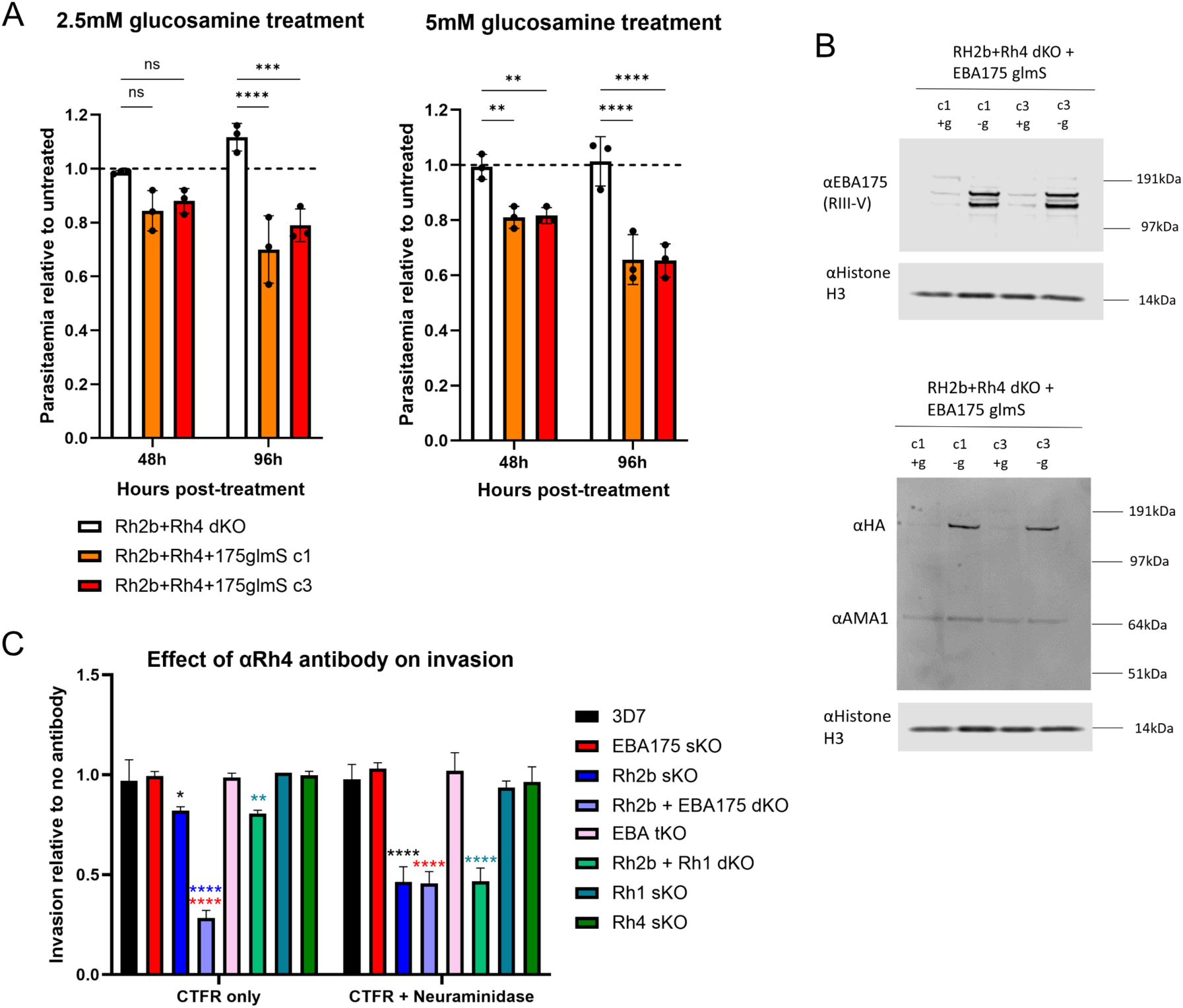
Disruption targeting PfRh2b, PfEBA175 and PfRh4 in combination causes an additive deficiency in invasion. A) 2.5mM or 5mM glucosamine was used to induce glmS ribozyme-mediated knockdown of PfEBA175 in the PfRh2b+PfRh4 double knockout background. The parasitaemia following induction with glucosamine for 48h or 96h is plotted as a proportion of the uninduced parasitaemia at the same time point. Two knockdown clones (PfRh2b+PfRh4+PfEBA175glmS c1 and c3) are shown with the PfRh2b+PfRh4 parental double knockout line included as a control (**P<0.01, ***P<0.001, ****P<0.0001, one-way analysis of variance with Dunnett’s multiple comparisons test). The error bars represent the mean ± s.d of three biological replicates. B) Western blots confirming glmS ribozyme-mediated knockdown of PfEBA175 following addition of 5mM glucosamine for 96h. PfEBA175 was detected with αEBA175 or αHA as the protein was tagged at the C-terminus with 3xHA. αHistone H3 was used as a loading control and αAMA1 was used to control for parasite stage. C) Parasite invasion into Cell Trace Far Red (CTFR) labelled red blood cells in the presence of αRh4 antibody relative to invasion in the absence of antibody. CTFR labelled cells were either untreated (CTFR only) or were pre-treated with neuraminidase (CTFR + Neuraminidase). Statistics are shown for comparisons between single knockout lines and 3D7 or double knockout cell lines and their corresponding single knockout lines (* are coloured to match the comparator line). Comparisons that did not pass the significance threshold (P<0.05) are not plotted (*P<0.05, **P<0.01, ****P<0.0001, one-way analysis of variance with Tukey’s multiple comparisons test). The error bars represent the mean ± s.d of three biological replicates.

As an alternative approach to interrogate the effect of disrupting PfEBA175, PfRh2b and PfRh4 in combination, the invasion capability of an PfRh2b+PfEBA175 double knockout cell line was assessed in the presence or absence of a monoclonal antibody targeting PfRh4 (Figure 5C). Invasion of the PfRh2b+PfEBA175 double knockout cell line into CTFR labelled RBCs was reduced by 70% in the presence of αRh4 antibody relative to the untreated control. There was a small but significant reduction in invasion by PfRh2b single knockout and PfRh2b+PfRh1 double knockout parasites in the presence of αRh4 antibody, whereas the invasion of PfEBA175 or PfRh1 single knockout and EBA triple knockout parasites was not impacted. Invasion into neuraminidase treated RBCs was also assessed in the presence or absence of αRh4 antibody. Neuraminidase treatment, to limit the use of EBA invasion ligands, in combination with αRh4 antibody treatment reduced the invasion of the PfRh2b single knockout and PfRh2b+PfRh1 double knockout cell lines significantly. For these PfRh2b knockout lines, this combination of treatments would be expected to impact all remaining EBA/Rh ligand-receptor interactions except for PfRh2a. The ability of these parasites to invade (albeit at a reduced capacity), indicates either an incomplete block by the αRh4 antibody or that residual PfRh2a activity may be sufficient for invasion. Overall, these results support a model where PfRh2b, PfRh4 and PfEBA175 are key invasion ligands, and disruption of these in combination causes an additive deficiency in invasion.

## Discussion

By combining a pooled transfection approach with Next Generation sequencing to detect edited parasites, we have developed a new scalable approach to testing the viability of combinatorial knockouts in *Plasmodium falciparum* parasites. We have applied this approach to systematically examine redundancy across the EBA and Rh invasion ligand families for the first time. This analysis indicated that all tested double knockout combinations were viable, significantly expanding on the few combined mutants that had been generated previously, and generating double mutants across the EBA/Rh families for the first time.

By employing amplicon sequencing to quantitatively compare the recovery of frameshift and synonymous edits in pools we were also able to identify mutant combinations that were viable but detrimental to parasite proliferation. However, while we are using the readout of frameshift edits in pooled transfections as an indication of combinatorial knockout viability, the recovery of frameshift edits could be impacted by various factors. Knockout combinations that cause an invasion defect but are still viable might be able to be generated in isolation but get outcompeted by other parasites within a pool. It is therefore critical to carry out independent transfections to confirm double mutant viability, as we did here. Editing efficiency of the specific gRNAs can also significantly influence recovery of edits, although we controlled for this by using the same gRNA to introduce frameshift and synonymous edits to the same genomic location. Nevertheless, there was still variation detectable in the pools, such as poor recovery of PfRh2a and Pfs230 frameshift edits in all pools, perhaps due to the quality of the transfected repair template DNA. Despite technical noise in the data, by focusing on consistent observations across the dataset, the pooled knockout approach proved valuable and allowed us to both more rapidly determine the full range of viable combinatorial mutants, and also identify combinations that were potentially detrimental, such as the PfRh2b+PfEBA175 mutant combination, which was validated in follow-up experiments.

Within the pooled analysis attempts to amplify the edited region of PfEBA140 in the PfRh2b single knockout background failed to produce an amplicon of the expected size. We therefore used Oxford Nanopore long read whole genome sequencing to generate whole genome sequences for the PfRh2b single knockout and PfRh2b+PfEBA175 double knockout lines, revealing complete loss of the end of chromosome 13 in a region that includes PfEBA140 as well as additional genes, primarily vars and rifins (Supplementary figure 8, Supplementary table 3). Spontaneous deletion of EBA and Rh genes has been observed before, particularly loss of PfRh2b in PfEBA181+PfEBA140 dKO parasites ^11^ and deletion of both PfEBA140 and PfRh2b genes in the D10 strain ^12,42^. We also carried out whole genome sequencing of the EBA triple KO cell line, which confirmed disruption of all EBA genes as expected, and also confirmed the presence of the preexisting mutations in the pseudogenes PfEBL1 and PfEBA165. However, once again we observed loss of coverage associated with a chromosome end, this time starting within the PfEBA181 gene (downstream of our FS edit) and extending towards the end of chromosome 1. In this case there appears to have been a rearrangement where a small section of chromosome 7 including the FS mutated PfEBA175 has been duplicated on chromosome 1 (Supplementary figure 9), though there remains no functional EBA sequence in this line. Genomic rearrangements are not uncommon in the sub-telomeres of *P. falciparum*^43^ and have been described in various field isolates, most notably deletion of the sub-telomere proximal hrp2 gene which encodes an antigen widely used in Rapid Diagnostic Tests^44^. It is possible that EBA genes, which are all located close to sub-telomeres, may be particularly prone to disruption through chromosomal rearrangements. These findings highlight that changes outside of the targeted genetic loci may arise during the course of generating clonal edited parasite lines, with genes close to the sub-telomeres particularly vulnerable. We suggest that the availability of relatively low-cost commercial long-read whole genome sequencing means that checks for genome rearrangement and telomere loss of critical lines should become a standard part of *P. falciparum* experimental genetic studies.

Our characterisation of double and triple mutants reveals several conclusions regarding long-standing questions about the interdependency and redundancy between EBA and Rh invasion ligands. Firstly, not all EBA ligands function equivalently. While loss of PfEBA175 altered invasion into enzyme treated cells and reduced parasite multiplication in shaking conditions, an EBA triple mutant that also lacked PfEBA140 and PfEBA181 behaved similarly to the PfEBA175 single mutant with no additive impact on invasion. PfEBA140 and PfEBA181 therefore do not compensate for the absence of PfEBA175 in the 3D7 line. Secondly, the fact that a triple knockout EBA line (PfEBA175+PfEBA140+PfEBA181) can be generated provides the first genetic evidence that *Plasmodium falciparum* can invade in the absence of a functional EBA ligand. This contrasts with *Plasmodium knowlesi* where loss of the EBA orthologue DBPα prevented invasion of human RBCs ^18^. Though loss of all *P. falciparum* EBA ligands is relatively well tolerated *in vitro*, the growth defect observed for a PfEBA175 mutant in shaking conditions suggests that under more challenging physiological conditions there would be a selective advantage to retaining their function. Of all the combinations tested, the PfRh2b+PfEBA175 and PfRh4+PfEBA175 double knockout cell lines had the most pronounced growth defects in both static and shaking conditions and these defects were stronger than seen in either the EBA triple knockout or in the PfRh2b+PfRh4 double knockout. This suggested that, as for the EBA proteins, not all Rh proteins are equivalent or interchangeable for efficient invasion, and that lines which combine mutations in both Rh and EBA families are more compromised, suggesting that the families could play different roles, a possibility discussed further below.

Our data clearly suggests that PfEBA175, PfRh2b and PfRh4 play dominant roles in invasion in the 3D7 line, consistent with previous observations ^38,42^. The effect of shaking on invasion of PfEBA175, PfRh2b, and PfRh4 single knockout lines was more pronounced than on the 3D7 parental strain, and the loss of each ligand individually altered the ability of the modified line to invade enzyme treated RBCs, pointing to a shift in receptor usage. While existing data from two lab strains has suggested that PfRh4 plays a role in compensation when EBA175 function is disrupted ^11,13^, characterisation of our knockout lines has revealed further detail about the compensatory interactions between these three key invasion ligands. The enzyme sensitivity profile of the PfRh2b+PfEBA175 double knockout line, alongside it’s sensitivity to inhibition by an αRh4 antibody supports the hypothesis that PfRh4 is compensating for the absence of these ligands. In contrast, the chymotrypsin sensitivity of the PfEBA175 single knockout was not a result of increased dependence on PfRh4 as the parasites did not become more sensitive to the αRh4 antibody. There is therefore potentially a molecular hierarchy where in the absence of PfEBA175 the parasites become more dependent on PfRh2b for invasion, but if PfRh2b is also disrupted there is a shift to dependence on PfRh4. PfRh4 function has previously been associated with efficient invasion in shaking conditions^45^. While our PfRh2b+PfEBA175 double knockout line is heavily dependent on PfRh4 for invasion it performs poorly in shaking conditions indicating that PfRh2b or PfEBA175 may also be important for invasion in shaking conditions, and that a combination of ligands cooperating to generate a stronger attachment could be more important in a dynamic moving environment.

Importantly, preference for the PfRh2b, PfRh4 and PfEBA175 invasion ligands may not be unique to lab strains such as 3D7. Data from Kenyan field isolates revealed either PfEBA175, PfRh2b or PfRh4 as the dominant transcript in various isolates and identified negative correlation between PfEBA175 and PfRh2b or PfRh4 transcript levels^15^. A similar trend was observed in Gambian isolates^16^. This consistency suggests the observations we have made here may be relevant to field strains. A previous study^11^ found that antibodies raised against PfEBA175, PfRh2a/Rh2b and PfRh4 in combination were more potent at inhibiting invasion than antibodies raised against the individual ligands. When an antibody is used to inhibit an invasion interaction the parasite has no time to adapt, if a targeted ligand is exposed on the parasite surface it can be bound by antibody. In contrast, our double mutants have had time to adapt to the absence of selected invasion ligands but nevertheless have been unable to recover the invasion efficiency of the parental 3D7 strain. This lends further support to the idea that an intervention strategy targeting PfEBA175, PfRh2b and PfRh4 in combination could be effective.

There are several possible models to explain the interconnected role of the EBA and Rh ligands in invasion. In one simple model, all EBA and Rh ligands can function equivalently in invasion, feeding into a single shared signalling pathway to support subsequent steps. In this mechanism some ligands could be superior to others through stronger interactions or more abundant receptors supporting a higher invasion rate. However, our data points to a more complex model, and does not support functional equivalence even within families. For instance, the PfEBA181 and PfEBA175 single knockout lines behave very differently in shaking conditions with the PfEBA181 mutant performing better and the PfEBA175 mutant worse.

In another model the EBA and Rh invasion ligands could function in successive interconnected steps. It has been proposed that PfRh1 may function in a signalling cascade that leads to PfEBA175 discharge from the micronemes, as monoclonal antibodies targeting PfRh1 inhibited this process^46^. However, the distinct invasion phenotypes of our PfRh2b+PfRh1 and PfRh2b+PfEBA175 double knockout lines suggest that the presence of PfRh1 may not be essential for PfEBA175 function in invasion, at least in the 3D7 line. Conversely, it has also been suggested that the engagement of PfEBA175 with its receptor Glycophorin A feeds into a signalling cascade that leads to the surface localisation of PfRh2b^47^. Again, given that an PfRh2b+PfEBA175 double knockout cell line had a stronger growth defect than a cell line lacking any functional EBA, it appears that EBA-receptor engagement is not essential for PfRh2b functionality *in vitro*. It is therefore difficult to reconcile our genetic data with a model in which the function of members of one gene family is strictly dependent on members of the other gene family, whichever the directionality of the dependence, e.g. Rhs depending on EBAs, or vice-versa.

A further possible model is that the EBA and Rh invasion ligands feed into separate and distinct pathways during engagement with the host erythrocyte, which promote successful invasion in different but complementary ways. This could involve making the host RBC more permissive for entry through cytoskeletal alterations^48^, enabling merozoite attachment to the host erythrocyte to facilitate further interactions, and/or signalling within the merozoite to feed into subsequent invasive actions. In this mechanism, one family may not be essential in relatively unchallenging *in vitro* conditions, as we found for the EBA family and the viability of the EBA triple knockout. By contrast, disrupting members of both EBA and Rh families could have a synergistic effect, as has previously been observed when different steps of invasion were targeted in combination using antibodies^49–51^, for example combined targeting of Rh4 and Rh5. This is the model that best fits the data presented here, and the potential for EBA and Rh invasion ligands to feed into separate signalling pathways is supported by previous experiments which swapped the C-terminal cytoplasmic domain of PfRh4 with that of PfEBA175 and found a lack of complementation^52^. In *Plasmodium knowlesi*, there is clear evidence of distinct actions for its EBA and Rh orthologues^18^, though the function of these ligand families may vary across divergent *Plasmodium* lineages. A complementary role for the EBA and Rh ligand families is supported by our double knockout data, where the absence of PfRh2b and PfEBA175, or PfRh4 and PfEBA175 is more detrimental in static and shaking conditions than the combined absence of PfRh2b and PfRh4. This indicates that the loss of key ligands from each family has a synergistic effect on invasion perhaps by reducing invasion efficiency in multiple different ways.

## Materials and Methods

### Ethics statement

Ethical approval for the use of human blood was obtained from National Health Service (NHS) Cambridge South Research Ethics Committee (20/EE/0100), and the University of Cambridge Human Biology Research ethics committee (HBREC.2019.40), formal written consent was obtained at the point of collection and is held by NHS Blood and Transplant (NHSBT).

### Plasmodium falciparum culture

The 3D7 strain of *P. falciparum* was cultured in human erythrocytes (from NHSBT, Cambridge, UK) at 4% haematocrit in RPMI 1640 medium (31870, Gibco), supplemented with 5g l^-1^ Albumax II, 2g l^-1^ dextrose anhydrous EP, 5.96g l^-1^ HEPES, 0.3g l^-1^ Sodium Bicarbonate EP, 0.05g l^-1^ hypoxanthine, 1% v/v L-glutamine and 0.05% v/v gentamicin. Cultures were maintained at 37°C, in gassed incubators with 1% O_2_ and 3% CO_2._

### Generation of EBA/RH frameshift edited single knockout cell lines

Frameshift mutations were introduced into the coding region of each EBA/Rh gene using CRISPR/Cas9 gRNA targeting. To generate the synonymous repair templates, a 300-500bp section of the binding region of each target gene was recoded to retain the amino acid sequence, but to remove guide targeting sites and allow identification of edited parasites. For each gene the frameshift repair template was identical in sequence to the synonymous repair template, except for the addition of 4 frameshift insertions and 6 mutations to create extra stop codons, as well as a 30bp sequence that was uniquely recoded to distinguish the frameshift from synonymous edit. Recoded sequences were synthesised by Twist bioscience or Genewiz (Azenta Life Sciences). 5’ and 3’ homology arms ∼800bp in length were amplified from 3D7 genomic DNA. Amplified homology regions and recoded repair templates were inserted into the pCC1 plasmid (PlasmoGEM Sanger) between SpeI and EcoRI sites using Gibson assembly (NEB Hifi assembly kit). Cas9 and guide RNAs were encoded on a separate plasmid. Two guide plasmids were used in this study, pDC2-cam-cas9-U6-yDHODH (PlasmoGEM Sanger) which was modified to include cytosine deaminase for negative selection, and pDC2-coCas9-gRNA-hDHFR gen2 (kind gift of Marcus Lee) which was modified to include cytosine deaminase and to remove a HA tag from the Cas9. To insert target-specific guide sequences into these plasmids, a pair of complementary oligonucleotides were designed for each guide which when annealed had overhangs that allowed ligation between two non-identical BbsI-generated sites in the plasmid. Oligonucleotide primers and synthesised sequences used in this study are listed in Supplementary table 4.

For individual transfections, 60µg of repair plasmid and 30µg of guide plasmid were combined, ethanol precipitated and resuspended in 30µl cytomix (120mM KCl, 0.15mM CaCl_2_, 2mM EGTA, 5mM MgCl_2_, 10mM KH_2_PO_4_, 25mM HEPES, pH 7.6). For pooled transfections targeting 2 genes, 15µg of synonymous repair plasmid and 15µg of frameshift repair plasmid were combined with 15µg guide plasmid for each gene. Ring stage parasites of 3-5% parasitaemia were washed in cytomix, and 150µl packed infected RBCs were mixed with the DNA and 300µl cytomix before electroporation using a Biorad Gene Pulser Xcell (Exponential mode, 310V, 950µF, resistance ∞).

Drug selection was applied the following day, DSM1 was used to select for the first edits and in most cases WR99210 (WR, Sigma) was used to select for subsequent rounds. WR was applied at 2.5nM, resistance to this compound was provided by hDHFR expression from the transfected guide plasmid ^53^. DSMI was applied at 1.5µM, resistance to this compound was provided by yDHODH expression from the transfected guide plasmid ^54^.

### Generation of conditional EBA175 knockdown cell line

To conditionally knock down EBA175 using the glmS ribozyme system^55^, a selection-linked integration approach^41^ was used to introduce a C-terminal 3xHA tag and glmS ribozyme sequence at the end of the gene. 885bp of the 3’ coding region of EBA175 was inserted into the pSLI HA-2A-NeoR-glmS plasmid^56^ (kind gift of Paul Gilson) between BglII and PstI sites using Gibson assembly. *P. falciparum* parasites were transfected with 60µg plasmid and were initially selected with WR99210 as described above. WR99210 resistant parasites were subsequently treated with G418 (Sigma-Aldrich) to select for parasites that had integrated the plasmid and so expressed the neomycin resistance marker as previously described^41^.

### Western Blotting

Haemolysis with 0.1% saponin (in PBS) in the presence of protease inhibitor was used to enrich for parasite material. Infected RBCs were incubated with 0.1% saponin (in PBS) with protease inhibitor for 10 minutes on ice. The pellet was then washed with PBS containing protease inhibitor and the dry pellet was stored at -70°C. The saponin lysed pellet was resuspended in ∼20x pellet volume of water with protease inhibitor before quantification using the Pierce BCA Protein Assay Kit (Thermo Scientific).

7.5µg of each sample was mixed with LDS sample buffer (NuPAGE, Invitrogen) and sample reducing agent prior to gel loading. Protein samples were resolved on 4-12% Bis-Tris SDS-PAGE gels and blotted onto nitrocellulose membrane. Primary antibody dilutions were prepared in 5% milk and the membrane was incubated overnight. The rabbit αEBA175 (region III-V) antibody (kind gift of Alan Cowman) was used at 1:1000. The mouse αAMA1 antibody was used at 1:1000 (This reagent was obtained through BEI Resources, NIAID, NIH: Monoclonal Anti-Plasmodium falciparum Apical Membrane Antigen 1 (AMA1), Clone N4-1F6 (produced in vitro) MRA-481A, contributed by Carole A. Long). The rabbit αHistone H3 antibody (abcam) and mouse αPfEXP2 antibody (European Malaria Reagent Repository) were used at 1:2000 for loading controls. Secondary antibodies were diluted in 50% TBS and 50% Li-cor blocking buffer. Both anti-mouse (IRDye 680RD Goat anti-mouse IgG, Li-cor) and anti-rabbit (IRDye 800CW Goat anti-rabbit IgG, Li-cor) secondary antibodies were diluted 1:7500. Immunofluorescence was detected on the Li-cor Odyssey imaging system.

### Next generation sequencing

Parasite material was isolated from the pooled transfections, and EBA/Rh target regions were first amplified using primers external to the homology arms used for integration. The amplified product was gel extracted and purified to prevent carryover of unintegrated plasmid. This product was then used as a template to amplify a 160-190bp fragment containing the 30bp frameshift unique sequence or synonymous sequence, the primers used in this step also added an overhang for subsequent addition of Illumina index primers. In the final amplification step Illumina Unique Dual Indexing primers were used to add unique identifiers for all samples. Indexed amplicons were then purified using AMPure beads (1:1 ratio) and quantified individually using a QuantiT assay (Invitrogen). Purified products were also run on an agarose gel to confirm addition of indexes. 100ng of each sample was pooled, and the pool was requantified using the QuantiT kit and diluted to 8nM. The pooled sample was quality checked on an Agilent tapestation prior to sequencing, and was sequenced on a Novaseq6000 instrument with a 50% PhiX spike in. Sequencing was carried out by Cancer Research UK.

The quality of the sequencing data was checked using FastQC. Depth and coverage were calculated with samtools v1.21. The number of reads recovered for the synonymous or frameshift sequence were counted using the bartender tool (https://github.com/LaoZZZZZ/bartender-1.1). Read 1 data was analysed as it was of higher quality than the read 2 data. The bartender tool is designed to count barcodes in sequencing data and when provided with invariant flanking sequences and the expected length of a central variant region, reports on the variant sequences observed and the number of reads corresponding to each sequence. Only the reads that exactly matched the expected synonymous or frameshift sequence were counted and plotted.

Example code:

`bartender_extractor_com \`

`-f UDP0017r_1.fq \`

`-o UDP0017r_1_extracted \`

`-q\`

`-p TAGAC [30] AATAA \`

`-m 0 bartender_single_com \`

`–f UDP0017r_1_extracted_barcode.txt \`

`–o UDP0017r_1_barcode \`

`–d 1`

### Growth assay in static and shaking conditions

Parasite cultures were synchronised using 5% D-sorbitol to retain ring stage parasites and eliminate later stages ^57^. The parasites were allowed to recover for at least 2 hours and were then diluted to 0.5% parasitaemia at a 4% haematocrit in 30ml. This was divided across 6x 5ml wells in 2x 6-well plates. One plate was placed on an orbital shaker (Celltron from Infors HT), which has a 25mm shaking throw, set to 90rpm, and incubated shaking at 37°C in a gas-controlled incubator (1%O_2_, 3%CO_2_). The other plate was kept in static conditions in the same incubator. After 48 hours, when the parasites had been through another invasion cycle and were again at the ring stage, the parasitaemia was quantified by flow cytometry. 20ul from each well was taken into a round-bottom 96-well plate and mixed with 100ul PBS containing SYBR green I (Invitrogen) at a 1:5000 dilution, the plate was then incubated at 37°C for 45min. Stained cells were washed once with PBS before resuspension in 200ul PBS and detection on an Attune NxT acoustic focusing cytometer (Invitrogen). SYBR green was excited with a 488nm laser (BL1-A) and detected by a 530/30 filter. 100,000 events were recorded per sample. The parasitaemia was measured using FlowJo™ software. Forward and side scatter area (FSC-A, SSC-A) was initially used to gate for the approximate size of an RBC, then the FSC-A and forward scatter height plot was used to gate for singlets. Parasitized RBCs were then counted by gating by BL1-A intensity to identify SYBR green positive cells.

### Invasion assays with enzyme treatment

To measure invasion into enzyme treated RBCs, RBCs were first labelled with 4µM cell trace far red (CTFR) for identification by flow cytometry. After 2h CTFR staining (rotating at 37°C), labelled cells were washed 1x in complete media, then 2x in RPMI only. CTFR labelled cells were resuspended in RPMI alone or RPMI with either 66.7mU/ml of neuraminidase (Type III from *V. cholerae*, Sigma N7885) or 1mg/ml chymotrypsin (Chymotrypsin, Alpha, TLCK Treated, Worthington Biochemical) added. After 1 hour incubation (rotating at 37°C), treated cells were washed 3x with RPMI and resuspended in complete media at 2% haematocrit. 60µl CTFR labelled RBCs were mixed with 60µl donor parasitized RBCs (1% parasitaemia, 2% haematocrit) in a 96 well round-bottomed plate to give a starting parasitaemia of 0.5% for all samples. After 48 hours, when the parasites had been through another invasion cycle and were again at the ring stage, the parasitaemia was quantified by flow cytometry (as described above). 100,000 events were recorded per sample. An additional gating step was included in the analysis of this assay, CTFR was excited with the 637nm laser (RL1-A) and detected by a 670/14 filter. Cells were gated as CTFR positive based on RL1-A intensity, and the proportion of parasitized RBCs (SYBR green positive cells) was calculated for this subset of cells. Invasion into enzyme treated cells was compared to invasion into untreated CTFR labelled cells.

### Invasion assays with αRh4 antibody

RBCs were labelled with CTFR and were then treated with neuraminidase or left untreated (as described above). 50µl of CTFR labelled target cells at 2% haematocrit were plated in wells of a 96 well round-bottomed plate. 5µl of αRh4 monoclonal antibody H12 ^58^ (kind gift from Wai-Hong Tham) was added to antibody treatment wells and 5µl of PBS to control wells. Finally, 50µl donor parasitized RBCs (1% parasitaemia, 2% haematocrit) were added to all wells. After 48 hours, when the parasites had been through another invasion cycle and were again at the ring stage, the parasitaemia in CTFR labelled cells was quantified by flow cytometry (as described above). Invasion in the presence of antibody was compared to invasion without antibody.

### Whole genome long-read sequencing (Oxford Nanopore Technology)

Haemolysis with 0.1% saponin (in PBS) on ice was used to enrich for parasite material. The pellet of parasite material was washed until the supernatant was clear and resuspended in 200µl PBS. 20µl proteinase K and 4µl RNase A (100mg/ml in PBS) were added and the sample processed using the DNeasy Blood and Tissue kit (Qiagen), finally eluting in 50µl buffer AE (10mM Tris-Cl, 0.5mM EDTA pH9). DNA purity ratios were checked on a NanoDrop spectrophotometer and quantified using a QuantiT assay (Invitrogen). To check samples were of sufficient fragment size for long-read sequencing they were run alongside the GeneRuler High Range DNA ladder (Thermo Scientific) on a 0.4% agarose gel. Quality checked samples were then submitted to Plasmidsaurus for long-read whole genome sequencing using Oxford Nanopore Technology. Plasmidsaurus assembled consensus contigs and provided raw reads. We aligned raw reads to the 3D7 reference genome downloaded from NCBI (Genome assembly GCA_000002765) using minimap2 (https://github.com/lh3/minimap2). Samtools was then used to create bam files which were used to visualise read alignments in IGV ^59^. Structural variation was further analysed using the SAVANA algorithm^60^.

## Author contributions

J.C.R. conceived the study, obtained funding and provided overall supervision. E.S. designed, performed and analysed the experiments, with J.C.R. and E.K. contributing to experimental assay design and interpretation. C.B. A.C. and A.K. provided experimental support. V.F. contributed to analysis of the sequencing data. E.S. and J.C.R. wrote the manuscript, E.K. assisted with editing the manuscript. All authors read and approved the manuscript.

## Acknowledgements

The authors would like to thank Kathryn Crouch for advice on analysis of next generation sequencing data. Wai-Hong Tham provided anti-Rh4 antibodies and Alan Cowman provided anti-EBA175 antibodies. Anti-AMA1 antibodies were obtained through BEI Resources, NIAID, NIH. Anti-EXP2 antibodies were obtained from the European Malaria Reagent Repository (EMRR: www.malariaresearch.eu). Marcus Lee provided the pDC2-coCas9-gRNA-hDHFR gen2 plasmid and Paul Gilson provided the pSLI HA-2A-NeoR-glmS plasmid. Ross F. Waller provided technical resources for analysis of sequencing data. This work was supported by the Wellcome Trust, United Kingdom, Investigator Award 220266/Z/20/Z to J.C.R. V.F. was funded by a Gordon and Betty Moore Foundation Investigator Award to Ross F. Waller (https://doi.org/10.37807/GBMF9194).

## Data availability

The short-read amplicon sequencing data and the whole genome long-read sequencing data have been deposited in the European Nucleotide Archive (ENA) under project number PRJEB107322.

## Declaration of interests

The authors declare no competing interests.

---

**Supplementary figure 1.**
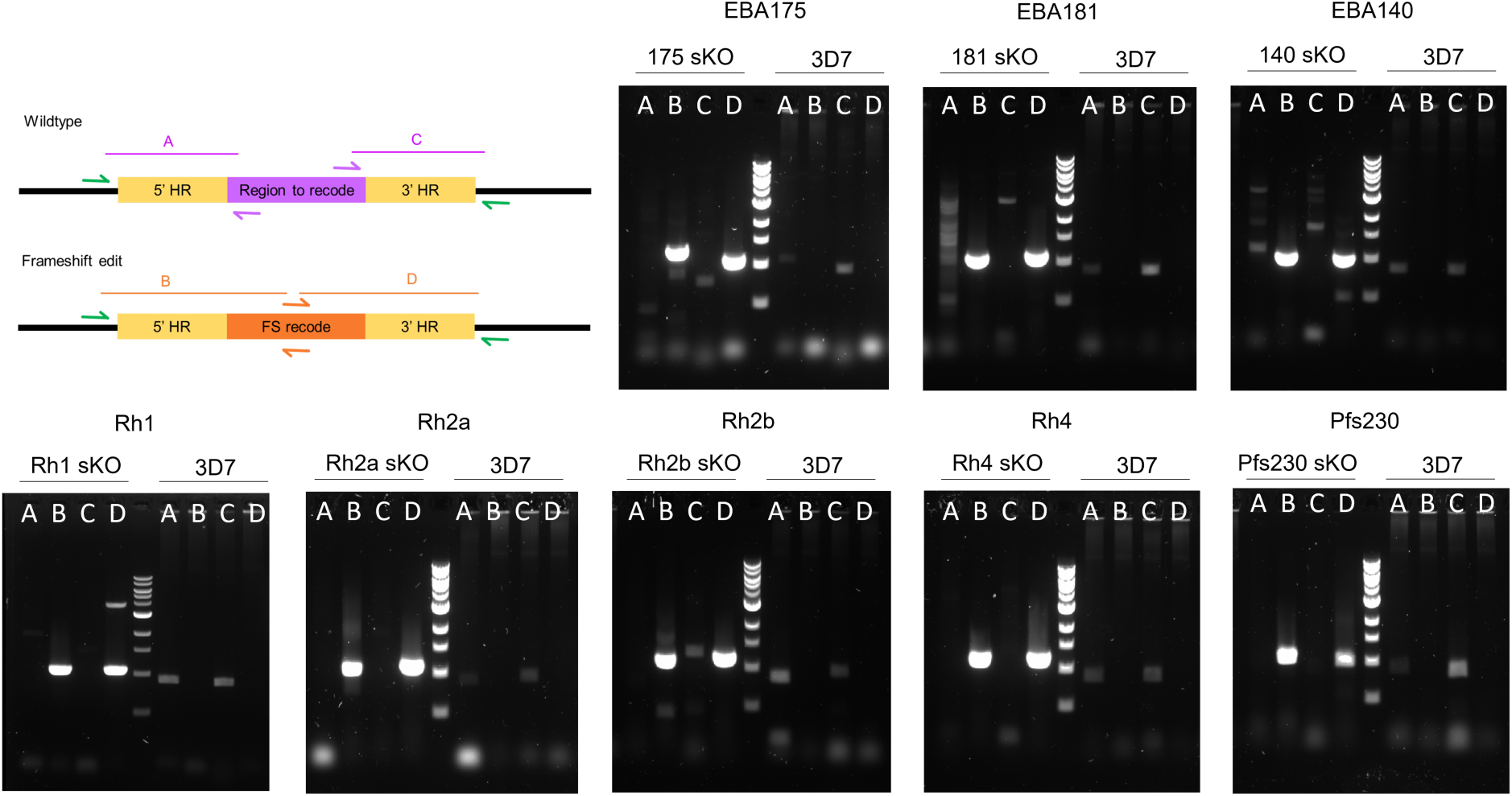
Genotyping PCRs confirmed integration of frameshift edit into EBA/Rh target loci. Reactions A and C are only expected to produce a product from the unedited wildtype locus. Reactions B and D are only expected to produce a product from the frameshift edited locus due to recoded sections of the repair template.

**Supplementary Figure 2.**
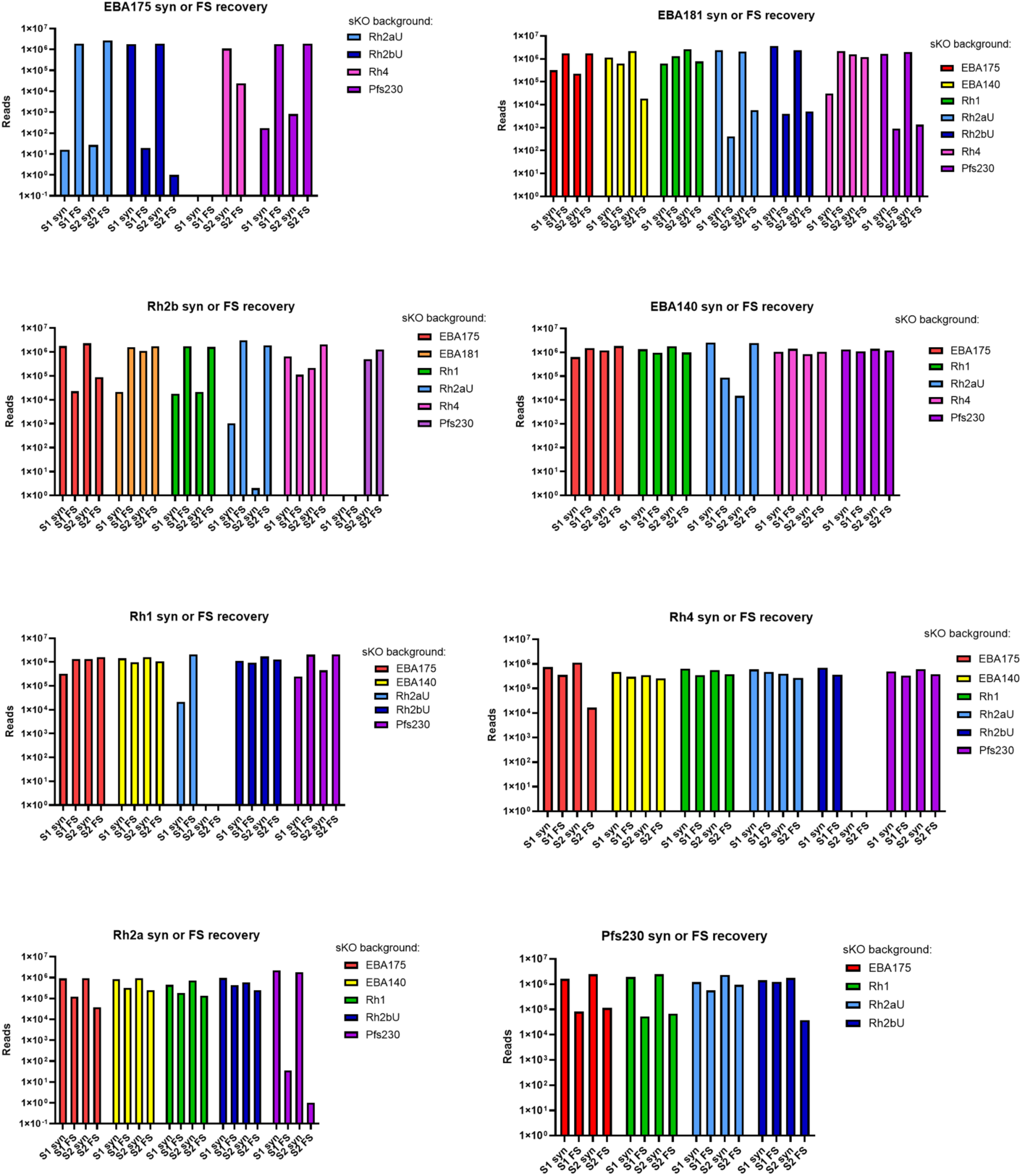
Next generation sequencing was used to detect synonymous and frameshift edits in targeted EBA and Rh genes. The total number of detected reads corresponding to a synonymous (syn) or frameshift (FS) edit are plotted for each gene on a separate graph, with the transfected cell lines these edits were detected in indicated by the colours in the legend. Samples were collected on 2 separate days for each pool and the reads detected in these 2 samples are plotted separately (S1 and S2).

**Supplementary Figure 3.**
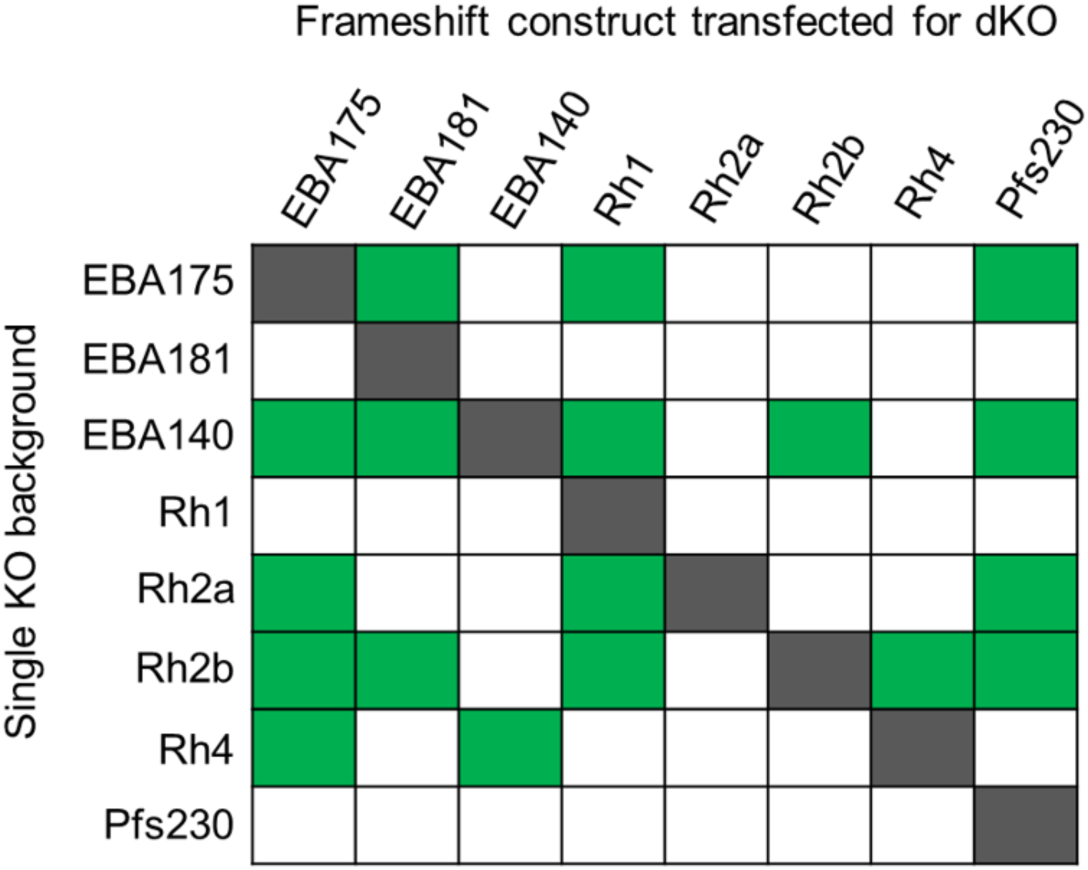
EBA and Rh double knockout combinations generated individually (highlighted in green).

**Supplementary Figure 4.**
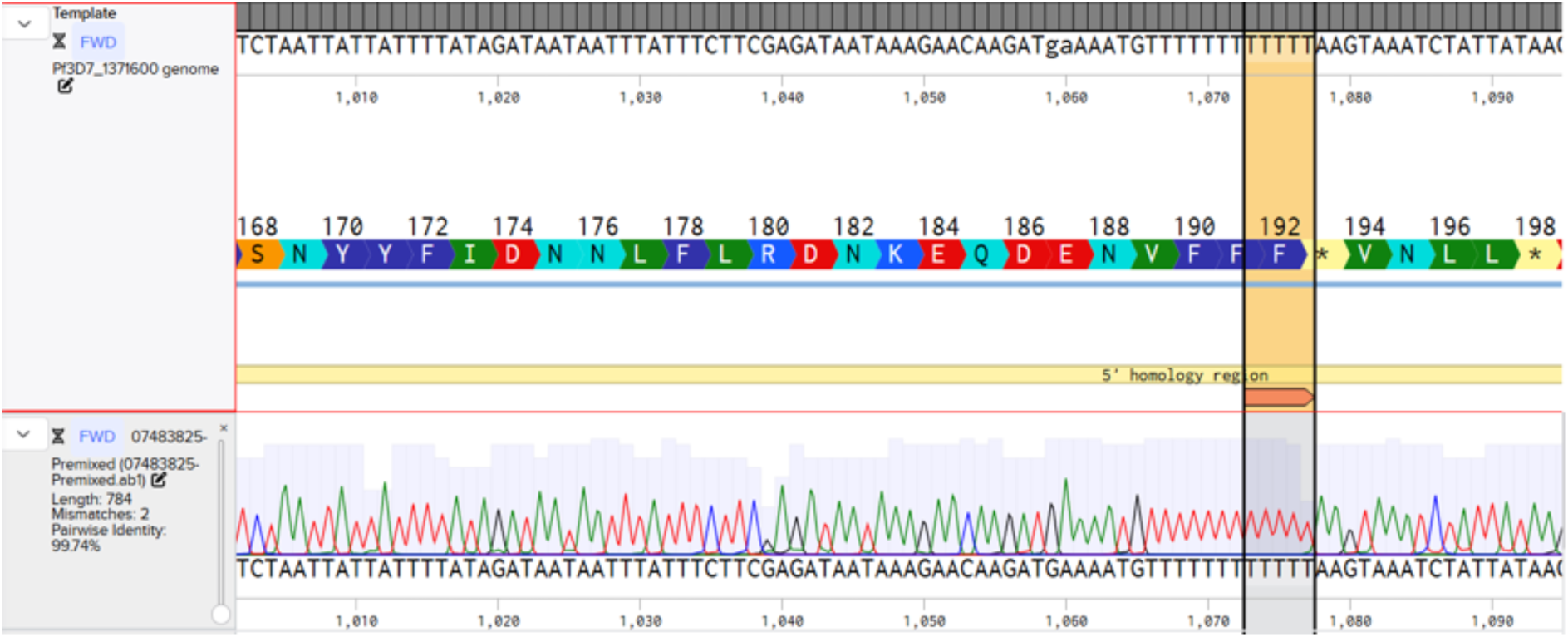
A 5T insertion is found in the coding region of EBL1 in the 3D7 strain, and the resultant frameshift introduces premature stop codons upstream of the DBL binding domain, this is expected to prevent expression of a functional EBL1 protein. Sequencing of the EBL1 locus from the EBA triple knockout cell line generated in this study revealed that this frameshift mutation was preserved. The figure shows the EBL1 sequence in the 3D7 strain (top row) aligned with the EBL1 sequence from the EBA triple knockout (bottom row) with the 5T frameshift insertion highlighted. This alignment was performed in Benchling [Biology Software]. (2025). Retrieved from https://benchling.com.

**Supplementary Figure 5.**
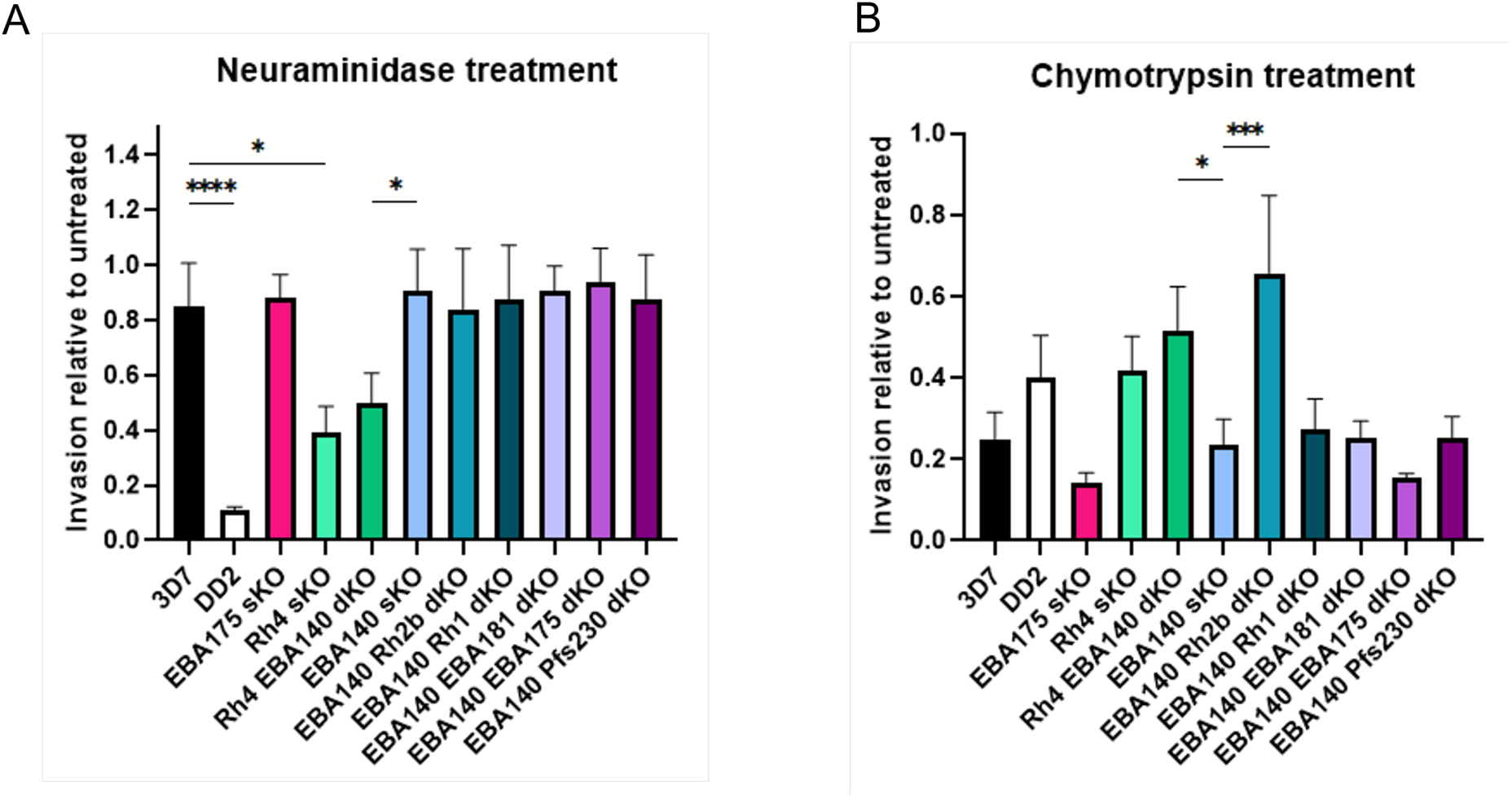
A) Parasite invasion into neuraminidase treated red blood cells relative to invasion into untreated red blood cells for EBA/Rh single and double knockout lines. B) Parasite invasion into chymotrypsin treated red blood cells relative to invasion into untreated red blood cells for EBA/Rh single and double knockout lines (*P<0.05, **P<0.01, ***P<0.001, ****P<0.0001, one-way analysis of variance with Tukey’s multiple comparisons test). The error bars represent the mean ± s.d of three biological replicates. Statistics are shown for comparisons between single knockout lines and 3D7 or double knockout cell lines and their corresponding single knockout lines. Comparisons that did not pass the significance threshold (P<0.05) are not plotted.

**Supplementary Figure 6.**
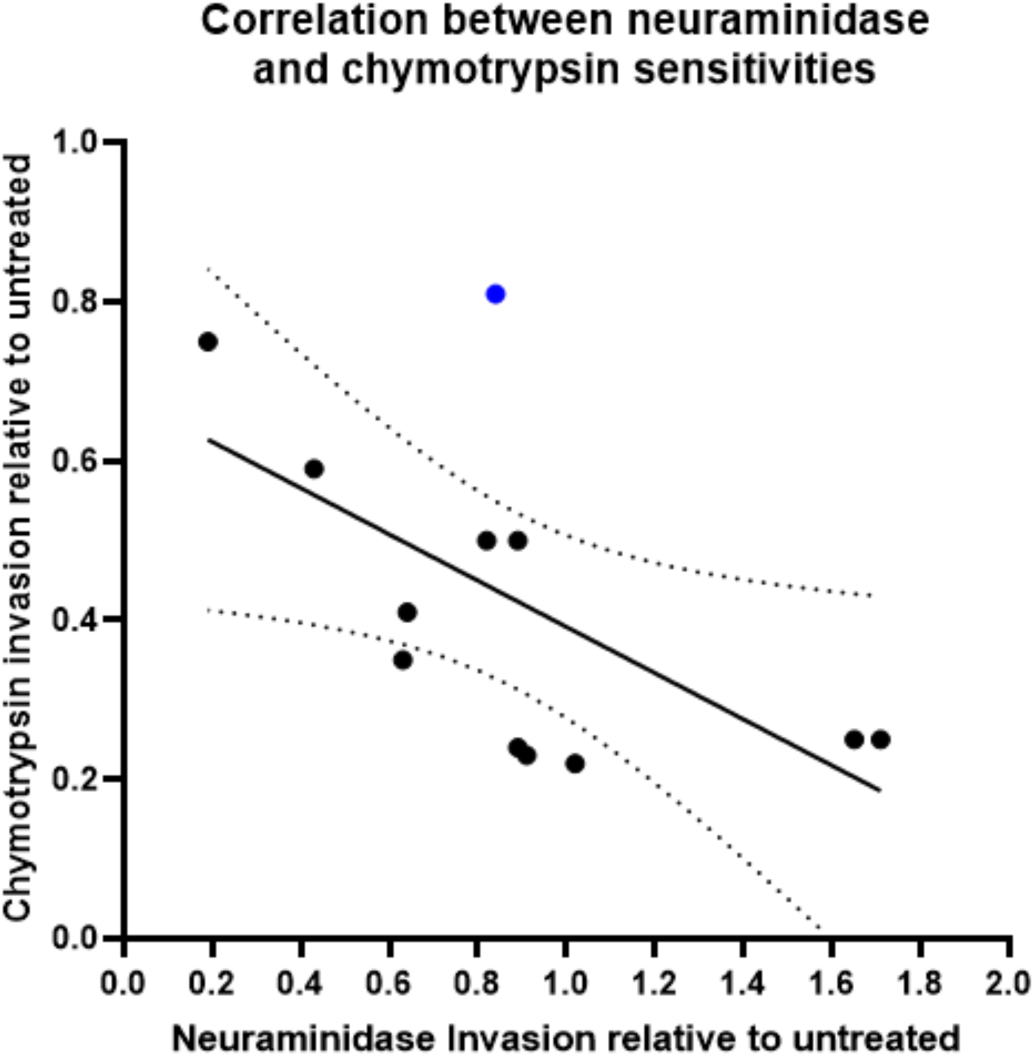
Simple linear regression revealing the negative correlation between neuraminidase and chymotrypsin sensitivities for EBA/Rh single and double knockout cell lines. Each data point represents the mean of three biological replicates. Highlighted in blue is the data point for the Rh2b single knockout, which is an outlier to the trend demonstrating resistance to chymotrypsin without a corresponding increase in neuraminidase sensitivity.

**Supplementary Figure 7.**
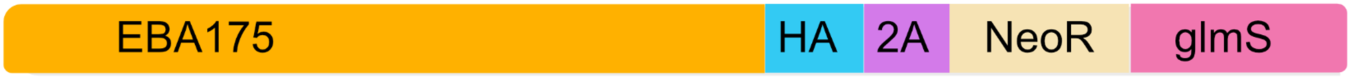
GlmS ribozyme addition to EBA175. A 3xHA tag was introduced at the C-terminus of EBA175, followed by a 2A skip peptide and neomycin resistance cassette. The 2A skip peptide allows separate expression of the EBA175 and neomycin resistance proteins from a shared transcript. The glmS ribozyme was inserted upstream of the 3’UTR to enable destabilisation of the transcript when ribozyme cleavage is activated by glucosamine addition.

**Supplementary Figure 8.**
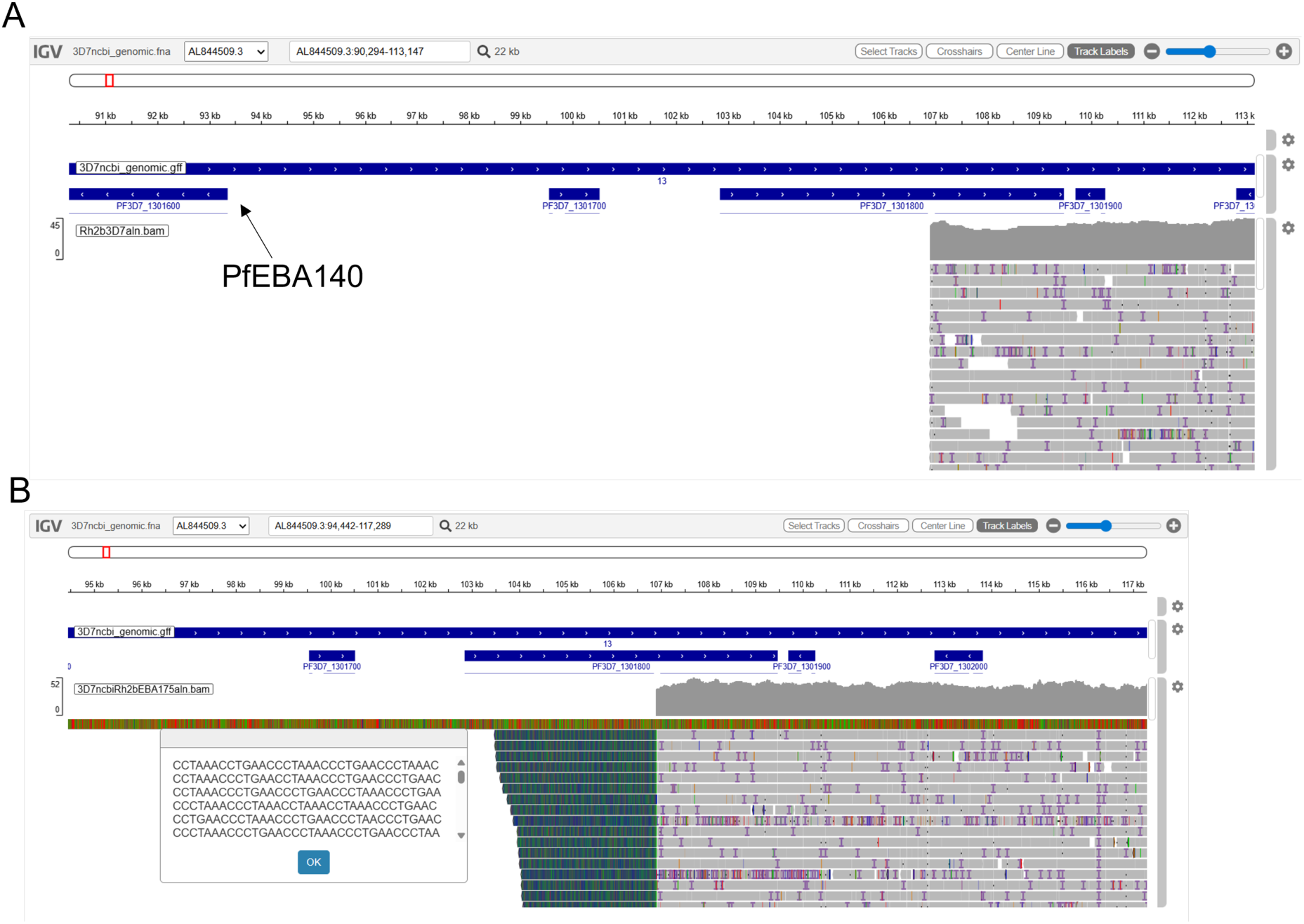
A) Visualisation of ONT reads derived from the PfRh2b single KO line aligned to the 3D7 reference genome on chromosome 13. The drop in coverage includes the PfEBA140 gene suggesting it has been lost in this line. B) Viewing the soft-clipped part of the reads covering the potential breakpoint reveal a sequence corresponding to telomere associated repeats (reverse complement) indicating that there was a break followed by telomere repair.

**Supplementary Figure 9.**
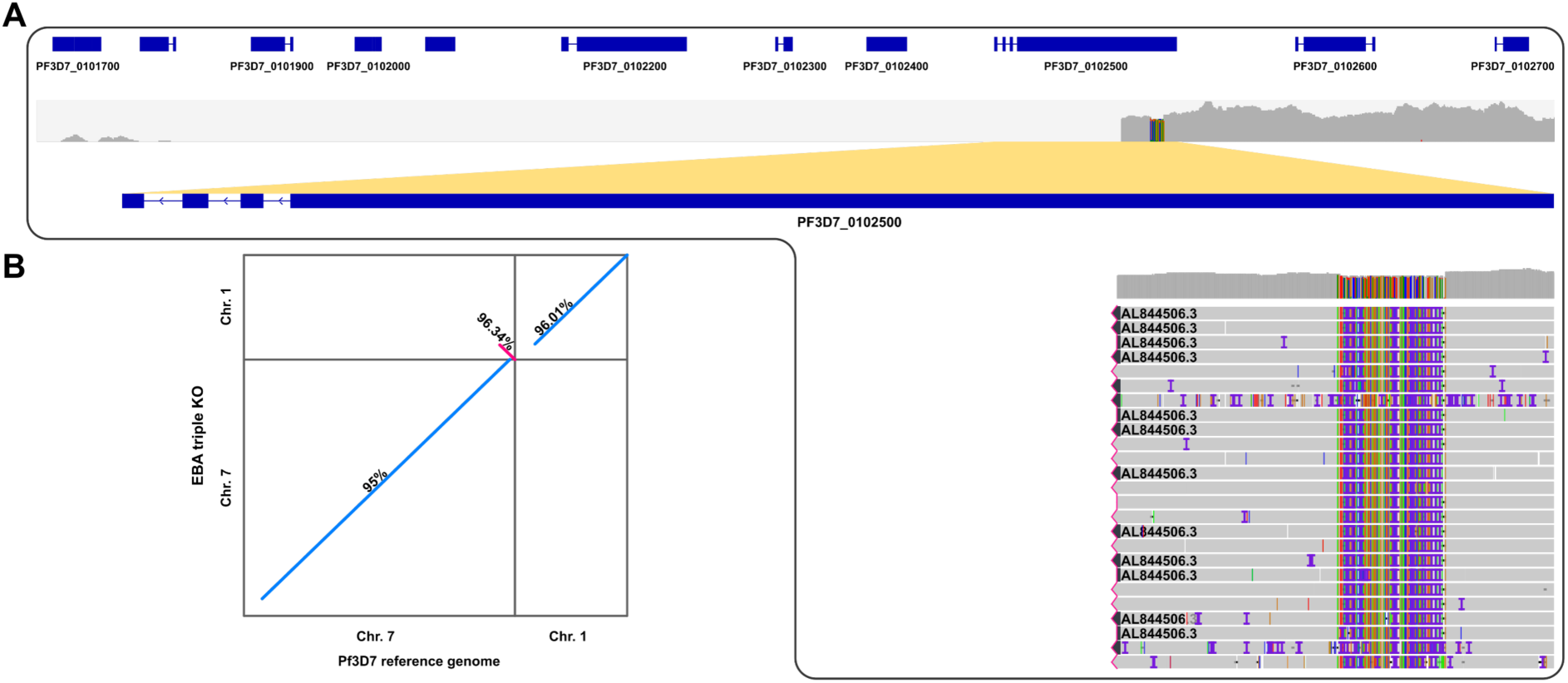
A) Visualisation of ONT reads derived from the EBA triple KO line aligned to the 3D7 reference genome on chromosome 1. The drop in coverage is located in the PfEBA181 gene, reads with secondary alignments indicate there has been a rearrangement with chromosome 7 (AL844506.3) B) Alignment of chromosomes 1 and 7 from the 3D7 reference genome and the assembled chromosomes of EBA triple knockout parasite line. Percent identities between sequences are shown next to the corresponding alignments. This alignment indicates there has been a rearrangement resulting in part of chromosome 7 being duplicated and translocated to chromosome 1. This structural variation was further confirmed by using SAVANA v1.3.5 (60).

## Notes

### Competing Interest Statement

The authors have declared no competing interest.

